# In vivo guanine quadruplex structure dynamics and role in genome maintenance in *Deinococcus radiodurans*

**DOI:** 10.1101/2025.05.27.656347

**Authors:** Himani Tewari, Shruti Mishra, Swathi Kota

## Abstract

Guanine quadruplex (G4) structures are secondary structures formed in nucleic acids that are rich in guanine bases. G4 structures regulate various cellular processes both in eukaryotes and prokaryotes. *Deinococcus radiodurans*, an extremophile show sensitivity to ionizing radiation in the presence of G4 binding ligands during the post-irradiation recovery period (PIR). Putative G4 motifs positioned at different locations on the genes fold into different topologies in vitro in the presence of monovalent cations while divalent cation Mg^+2^ supported the stable G4 formation but Mn^+2^ destabilizes the G4 structures. Thioflavin T and anti-DNA quadruplex antibodies detected the in vivo formation and dynamics of G4 structure in response to various DNA damaging agents, more in the presence of gamma radiation treatment. G4 binding drugs during PIR arrested the G4 structure dynamics and delayed the DNA repair process. Further, the absence of G4 helicase, RecQ resulted in the accumulation of more G4 structures and genome instability. All these results indicate that the guanine quadruplex structure formation increases in response to cellular stress and G4s are crucial for stable genome maintenance in *Deinococcus radiodurans*.

## Introduction

Guanine quadruplex (G4) structures are non-canonical secondary structures formed in DNA and RNA containing guanine-rich sequences. Four guanine bases pair through Hoogsteen hydrogen bonding to form a G-quartet, which upon stacking on each other form G-quadruplex structures. Monovalent cations such as Na^+^ and K^+^ play a significant role in the folding and stability of these secondary structures (Bochman et al. 2012). The regulatory role of G-quadruplex structures in different cellular processes and genome maintenance is well studied in higher eukaryotes (Maizels and Gray, 2013). G4 DNA structures found at telomeres and in the promoter regions of proto-oncogenic like cMyc genes are also being explored as targets for cancer therapies (Kosiol et al. 2021). In bacteria, the association of G4 structures with cellular processes is less explored, although some very specific roles of quadruplexes have been described. The G4 motif was first reported in pilin antigenic variation in the human pathogen *Neisseria gonorrhoeae* (Cahoon and Seifert, 2009). In silico analysis identified 52 putative G-quadruplex forming motifs in *E.coli,* (Kaplan et al., 2016) and at a few TSS on the genome of *Mycobacterium tuberculosis* (Perrone et al., 2017). Further, a systemic study was conducted on the positional influence of G4s on gene expression in *E. coli* by inserting G4s into different locations (sense and antisense strands, promoter, 5’ UTR, and 3’ UTR) of a GFP gene on a plasmid (Holder and Hartig 2014).

*Deinococcus radiodurans* is a gram-positive bacterium, known for its resistance to many abiotic stresses including gamma radiation. Various mechanisms like efficient DNA damage repair pathways, strong anti-oxidant systems, multipartite genome, and high GC content contribute to the extreme phenotype of this bacterium (Slade and Radman, 2011; Misra et al., 2013). The multipartite genome harbors 2 chromosomes and two plasmids (White et al., 1999). Numerous putative guanine quadruplex structure forming motifs (PQS) were identified across the GC rich genome. Earlier, it was shown that the presence of G4 stabilizing ligands like NMM (*N*-methyl mesoporphyrin IX) during the PIR makes the bacterium sensitive to ionizing radiation stress. The PQS in the upstream region of DNA repair genes like *recA*, *recQ*, *mutL,* etc. form stable G4 structures *in vitro* and regulate the gene expression (Kota et al., 2015). High throughput transcriptome analysis of *D. radiodurans* cells as a function of G4 stabilizing ligand NMM, and gamma radiation has also shown the role of G4 DNA motifs in the regulation of DNA damage responsive gene expression during PIR (Mishra et al., 2019). However, in vivo G4s formation and their function during the DNA damage recovery period is not known fully. Studies demonstrating the in vivo dynamics of G4 structures using fluorescent G4 specific dyes or antibodies are abundant in eukaryotes, but there exists very less or no studies in prokaryotes, that may unveil the role of these structures in phenotypes like radiation resistance or pathogenicity.

This study has investigated the in vivo formation of G4 structures and their effects on genome maintenance in *D. radiodurans*. The PQS present at different positions of the gene (5’, 3’, or intergenic regions) folds into different topologies in the presence of various cations. The presence of divalent cation Mg2+ supported the stable G4s formation, but in the presence of Mn^2+^, the G4s are not formed. Thioflavin T, a fluorescent dye and antibodies detected in vivo G4 structures and their dynamics during the PIR. Further studies showed that the arrest of G4 structure dynamics slows the DNA damage repair process and the absence of G4 helicase RecQ results in genome instability. All these results suggest that the in vivo G4 structure dynamics during the DNA damage recovery phase in *D. radiodurans* may play a crucial role in radioresistance and genome maintenance.

## 2. Materials and Methods

### 2.1 Bacterial strains, and materials

*Deinococcus radiodurans* R1 (ATCC 13939, a gift from J. Ortner, Germany, Schaefer et al. 2000), was grown at 32^0^C in TGY medium (Bacto Tryptone [0.5%], glucose [0.1%] and Bacto yeast extract [0.3%]). The *recQ* mutant (*ΔrecQ*) (Khairnar et al. 2019) of *D. radiodurans* cells was also grown overnight at 32°C in kanamycin (8 µg/ml) containing TGY medium. Anti-DNA G-quadruplex (G4) antibody clone 1H6 (MABE1126), 5,10,15,20-tetrakis-(*N*-methyl-4-pyridyl) porphyrin (TmPyP4), and 2-(4-(Dimethylamino)phenyl)-3,6-dimethylbenzo[d]thiazol-3-ium chloride (ThioflavinT, ThT dye) were procured from Sigma-Aldrich, St. Louis, USA. N-methyl mesoporphyrin (NMM) was obtained from Frontier Scientific, UT, USA, and all the chemicals and enzymes for molecular biology were procured from Merck Inc. and New England Biolabs. The molecular biology techniques used in this study were used as described earlier (Sambrook and Russell 2001).

### 2.2 Identification of putative DNA G-quadruplexes

The PQS present at different positions like 5’/3’ ends of the gene, coding regions, and in the intergenic regions on the genome of *Deinococcus radiodurans* were identified using QGRS (http://bioinformatics.ramapo.edu/QGRS/index.php) and G4 hunter(http://bioinformatics.ibp.cz/) software. For this, the FASTA format of chromosomes I and II along with megaplasmid and small plasmid were retrieved from NCBI and analyzed. The 3G run PQS present at different regions of the gene were selected and characterized further as described earlier (Table 1; Kota et al. 2015).

**Table 1.**
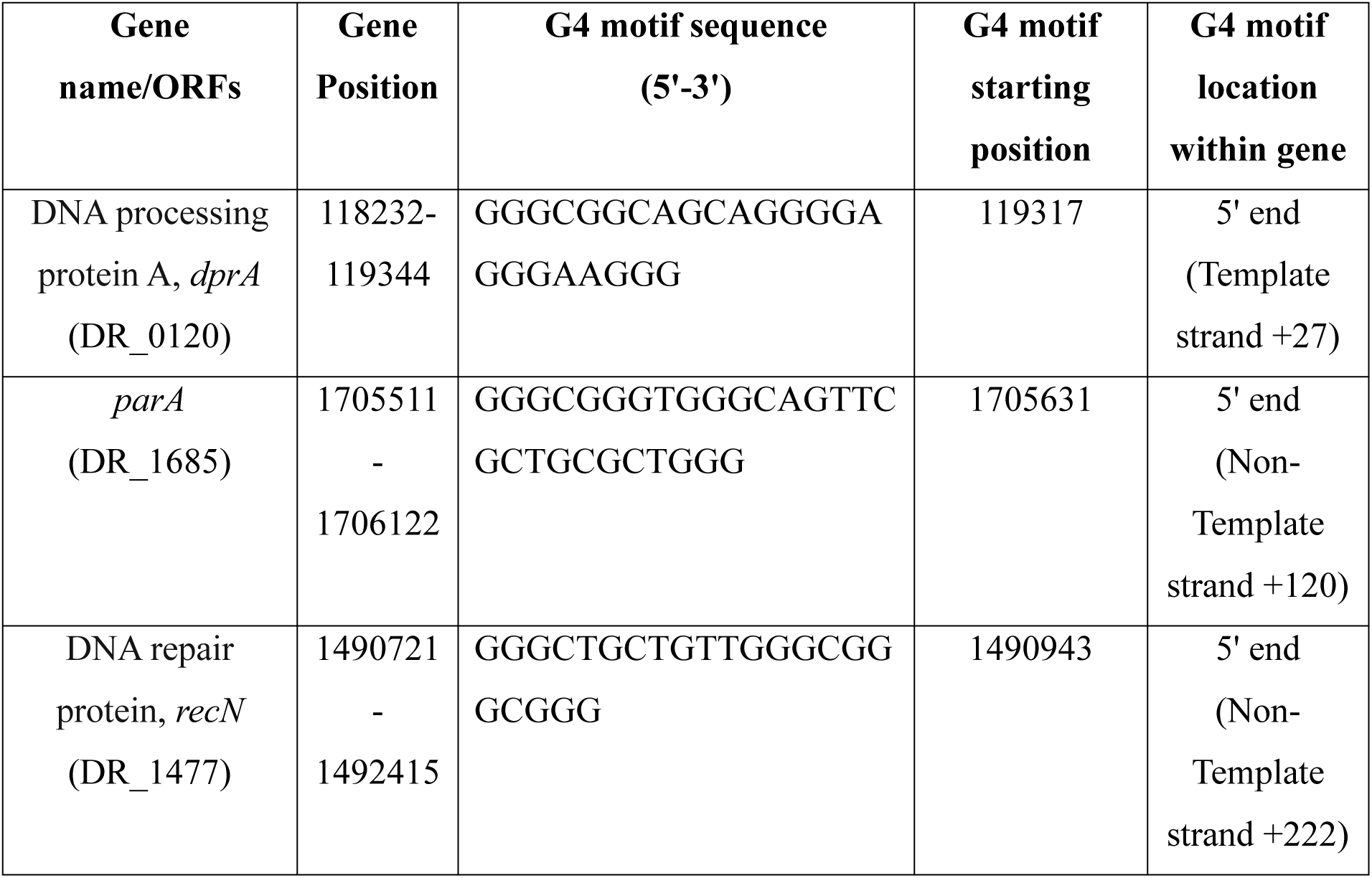
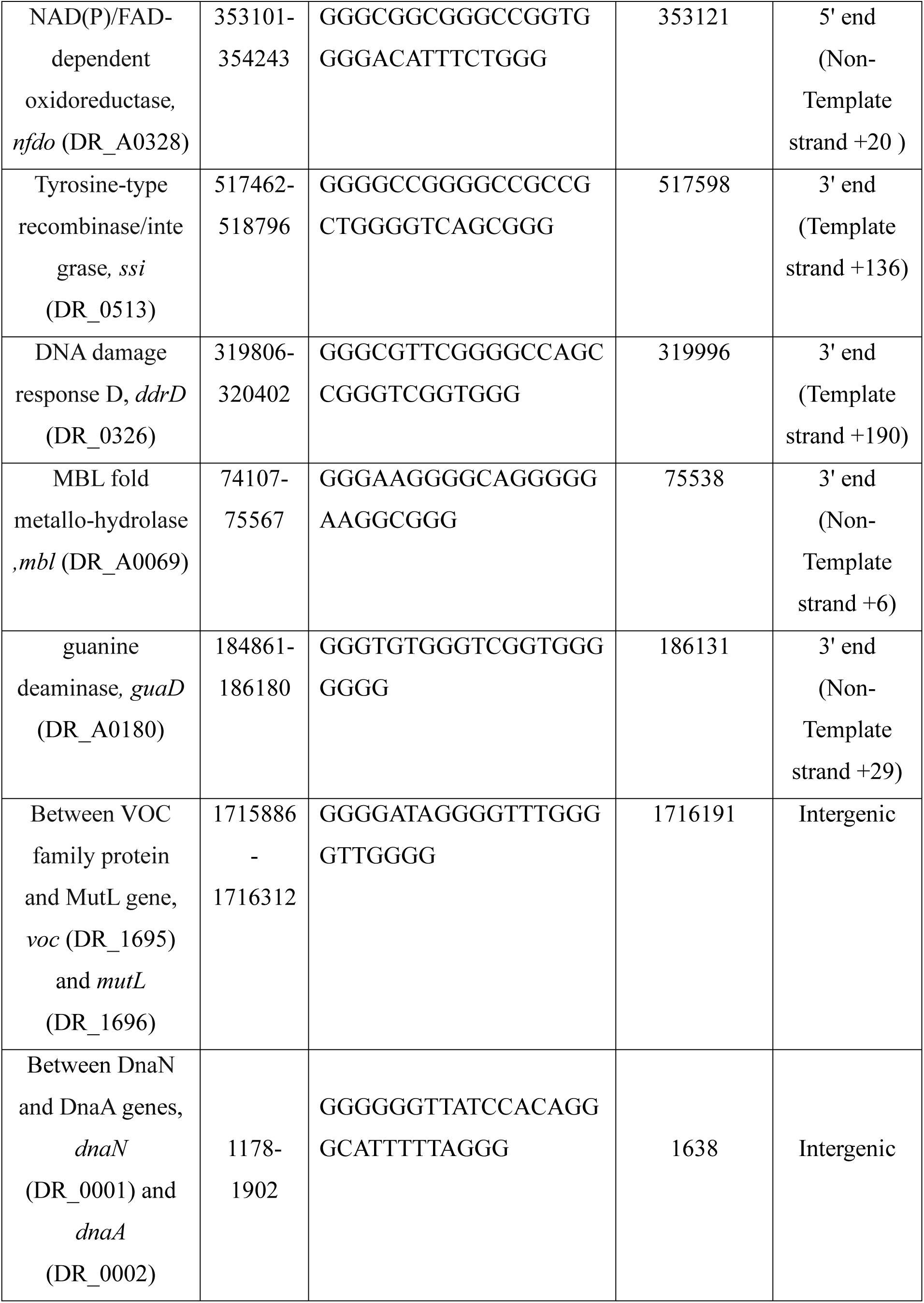
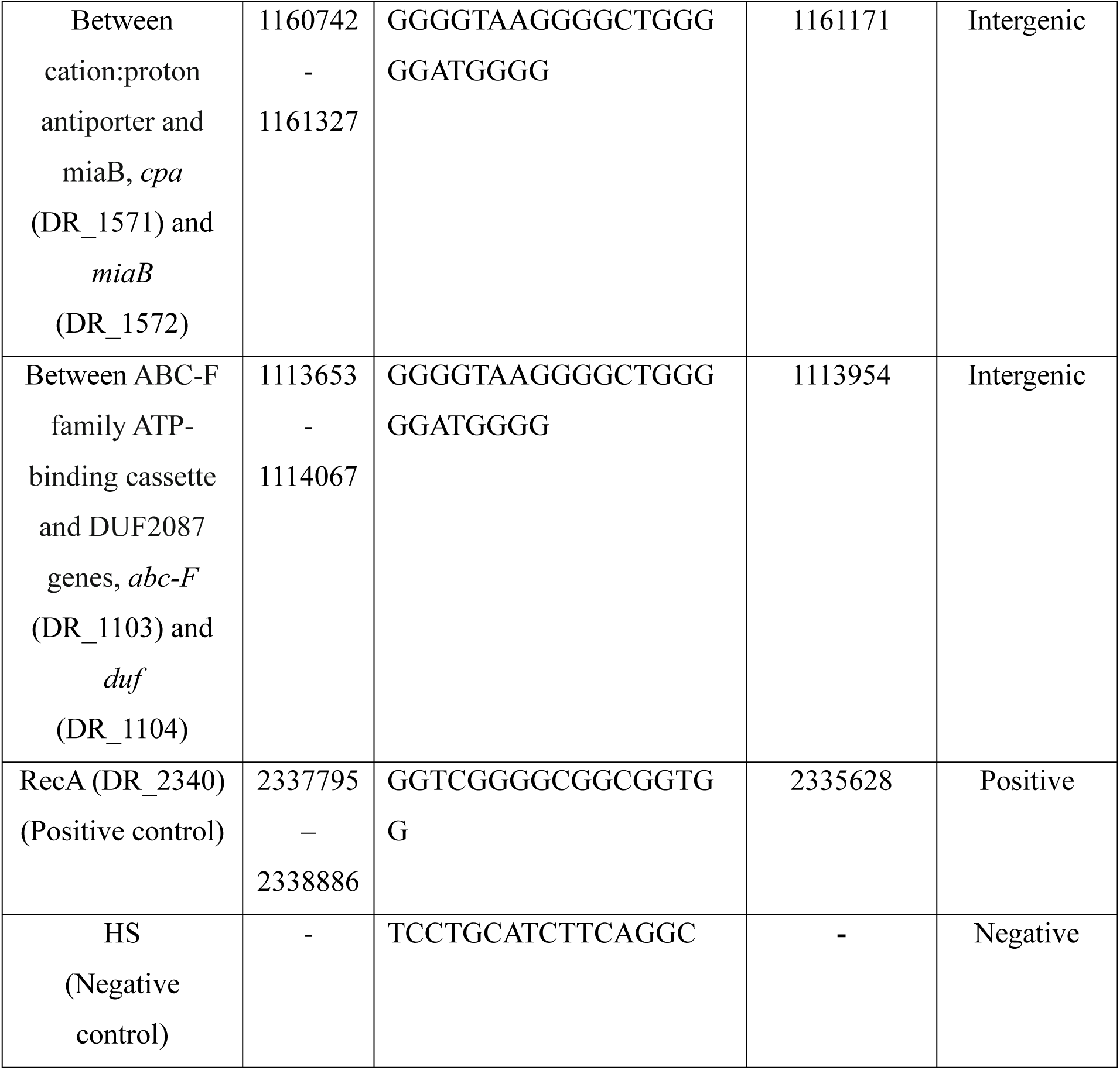
Putative G-quadruplex forming structural motifs in the genome of *Deinococcus radiodurans* used in this study.

### 2.3. Validation of in vitro G4 structure formation in the presence of various salts

PQS were checked for G4 structure formation in the presence of monovalent cations KCl (50 mM, 100mM), NaCl (50 mM), LiCl (50 mM), or divalent cations MgCl_2_ (10 mM) and MnCl_2_ (0.1, 2 and 10 mM) along with 10 mM Tris buffer (pH 7.4) by annealing the DNA sequences (5 μM) at 95 °C for 10 min and slowly cool down to room temperature. For the detection of G4 structure formation thioflavin T (ThT) fluorescence enhancement assays were carried out. For this, the annealed sequences were mixed with 0.5μM Thioflavin T (ThT) in a 96-well microplate. The fluorescence spectra were collected between 440 and 700nm after excitation at 425nm in Synergy H1 Hybrid multi-mode microplate reader. The fluorescence intensity of the ThT alone or in the putative G4-forming sequences was compared. The fluorescence emission spectra of ThT were plotted using GraphPad Prism 8.0. The fold change in fluorescence enhancement was calculated as described earlier (Renaud et al. 2014).

The topology attained by various PQS in the presence of different cations was examined using circular dichroism (CD) spectroscopy (Borgognoone et al. 2010). The CD spectra of the annealed sequences were recorded on a spectrophotometer (Biologic MOS-500 CD spectropolarimeter), in the wavelength range from 220 to 320 nm with a scanning speed of 100 nm/min and a response time of 2s, using a quartz cuvette with a path length of 1.0-mm. A non-G4-forming DNA sequence as a negative control (HS) and a known G4-forming sequence as a positive control (RecA) were taken in addition to putative G4-forming sequences (Mishra et al. 2019).

### 2.4 Cell survival studies

The exponential phase of *D. radiodurans* R1 cells grown at 32 °C in the presence and absence of 100 nM of NMM in TYG medium were exposed to 6 kGy γ radiation having a dose rate of 3.75 kGy/h (Gamma Cell 5000,^60^Co., Board of Radiation and Isotopes Technology, DAE, Mumbai, India). Cells were exposed to other DNA damaging agents like, H_2_O_2_ (70 mM and 100 mM), UV rays (1000 J/m^2^), and mitomycin C (MMC-5µM) in the presence and absence of NMM. After treatment, the cells were pelleted, and diluted to 20-fold in fresh TYG medium, with or without NMM, and growth was monitored at 600 nm in a Synergy H1 multimode plate reader for 18h. The growth curve data obtained was analyzed using GraphPad Prism 8.0 software and plotted.

### 2.5 Microscopic studies

Invivo formation of G4 structures was monitored by staining with ThioflavinT. The *D. radiodurans* R1 cells were grown overnight at 32 °C in TGY medium and exposed to various DNA damaging agents as discussed earlier. After treatment, the cells were allowed to recover with constant shaking, and collected at 1h and 4 h post-treatment. For G4 detection by Thioflavin T, the cells were washed in PBS and stained with 2 μg ml^-1^ DAPI (4′,6-diamidino-2-phenylindole, Dihydrochloride) for nucleoid staining and 2μM ThT (Thioflavin T) for G-quadruplex visualization. These cells were washed in PBS to remove excess dye and resuspended in a small volume of PBS. Two to three microliters of cell suspension were dropped on a glass slide coated with 0.8% agarose and covered with glass coverslips. The confocal microscopy was performed using Olympus IX83 inverted microscope as described earlier (Mishra et al. 2022). The DAPI fluorescence was visualized using a filter having 460 nm emission when excited at 402 nm. ThT fluorescence was monitored at excitation wavelength 438 nm and emission was recorded at 560 nm using the FITC channel. Image analysis was carried out using the built-in CellSense software. The brightness and contrast were adjusted using Adobe Photoshop 7.0. Approximately 200-250 cells from three independent experiments were examined for quantitative analysis and data was plotted using Graphpad Prism 8.0.

The formation of G4 structure in vivo was also detected by immunofluorescence microscopy using an anti-DNA G-quadruplex (G4) antibody (clone 1H6) which specifically targets G4 structures as described earlier (Manguan et al. 2014). Briefly, *D. radiodurans* wild type (R1) treated and untreated with NMM were subjected to 6 kGy radiation and were recovered with fresh medium. Post-irradiation recovering samples were collected at regular time intervals. The cells were fixed with a 4% paraformaldehyde solution and then washed in PBS. Further, the cells were permeabilized by treatment with lysozyme (2 mg/ml) for 1h at 37^◦^C, then with 0.1% Triton X-100 in PBS at 37^◦^C for 5 min. The cells were washed with PBS, resuspended, and around 5µl was spread onto a poly-l-lysine pretreated slide and allowed to air dry. Fixation was done by treating the slides with 4% PFA at 37 ^◦^C for 30 min. Using a blocking solution of 2% BSA in PBS-T (0.05% Tween 20 in PBS), samples were blocked for 2 hours at 37°C and subsequently incubated with the G4 antibody overnight in the same solution. Cells were washed with PBS-T for 20 min twice and then incubated with a secondary antibody, anti-mouse IgG conjugated with Alexa flour 594 (Sigma) for 2 h in a blocking solution. Cells were again washed with PBS-T for 20 min twice and slides were mounted using a mounting medium containing antifade DAPI. Cells were visualized under a microscope and the fluorescent intensity profile of ∼200 cells was analyzed using automated Olympus CellSens software and plotted using GraphPad Prism 8 software.

For morphological studies, both the wild type and *ΔrecQ* cells were grown, washed with PBS, stained with DAPI, and visualized. Approximately 200 cells were analyzed using Olympus CellSens software to quantify various cell morphologies. The resulting data was plotted with GraphPad Prism 8.0 software and differences were considered significant if the P-value was below 0.05 (95% confidence interval).

### 2.6 Pulsed-field gel electrophoresis (PFGE)

The repair of damaged DNA is monitored by pulsed-field gel electrophoresis, as described earlier (Mattimore and Batista 1996). Briefly, the cells were grown with and without NMM resuspended in fresh medium and were irradiated with a dose of 6 kGy. The cells were diluted in TGY and allowed for post-irradiation recovery and samples were collected at different time intervals. Each collected aliquot was pelleted by centrifugation and resuspended in butanol-saturated phosphate buffer followed by another centrifugation step. The obtained pellets were further resuspended in 0.5M EDTA, and kept at 60^0^C for 30 min afterward pelleted and resuspended in Tris-EDTA buffer. A cell suspension was mixed 1:1 with 1.5% low melting point agarose and solidified into plugs using a mold. These plugs were incubated overnight with lysis solution containing 1mg/ml of lysozyme at 37^0^C and later overnight incubated at 50 °C in 1 % sarcosine and 1 mg/ml proteinase K solution. The plugs were subjected to pulsed-field gel electrophoresis in 0.5XTBE using a CHEF-DR® III electrophoresis system (Bio-Rad) at 6 V/cm^2^ for 16 h at 14°C, with a linear pulse ramp of 10-60s and a switching angle of 120^0^. The gel was stained with Ethidium Bromide for visualization and documentation.

### 2.7 Statistical analysis

For all the statistical analysis Student’s t-test was used. The significance value-P value obtained at 95% confidence intervals are depicted as **** for P value <0.0001, *** for P value <0.001, ** for P value of 0.05-0.001, * for P value <0.05.

### 3.0 Results and Discussion

### 3.1 Putative guanine quadruplex structure-forming sequences fold into different topologies

Previously presence of PQS at the upstream regions of most of the DNA repair genes in *D. radiodurans* was reported (Kota et al. 2015). Here the existence of PQS at different positions to the genes - towards the 5’ or 3’ ends of the genes or intergenic regions was searched as described in the methodology. The analysis results showed the presence of 3 G run PQS at different locations on the genome. A few genes like - DNA repair protein (*recN)*, DNA processing protein A (*dprA)*, NAD(P)/FAD-dependent oxidoreductase (*nfdo*), and *parA* contain PQS towards their 5′ end. Similarly, genes like - DNA damage response D (*ddrD)*, tyrosine-type recombinase/integrase(*ssi*), guanine deaminase (*guaD),* and MBL fold metallo-hydrolase (*mbl*) contain towards 3′ ends. Few PQS were also found at intergenic regions-between VOC family protein and mutL, DnaN and DnaA, cation: proton antiporter and miaB and ABC-F family ATP-binding cassette and DUF2087 (Table 1). Further studies were carried out to check the G4 structure formation by PQS in the presence of different salts.

The formation of G4 structures by putative motifs was checked by ThT fluorescence assays as described in the methodology. It is known that Thioflavin T (ThT) is a benzothiazole that becomes fluorescent in the presence of the G4 structures (Renaud de la Faverie). The fluorescence intensity of the ThT alone or with the putative G4-forming sequences was compared. ThT exhibited an emission maximum at approximately 490 nm across all tested sequences, and only the presence of G4 DNA structures resulted in a rise in fluorescence. ThT fluorescence enhancement was observed in the positive control (*recA*) while no significant ThT enhancement was observed in the non-G4 sequence (HS), the signal obtained was similar to ThT alone, suggesting that these putative sequences may stably fold into G4 structure in vitro (Fig 1A). Further, the quantitative analysis showed that all the oligonucleotides containing PQS lead to an increase in ThT fluorescence (∼/> 5 fold increase in enhancement) compared to non-G4 sequence and ThT alone (Fig 1B).

**Figure 1:**
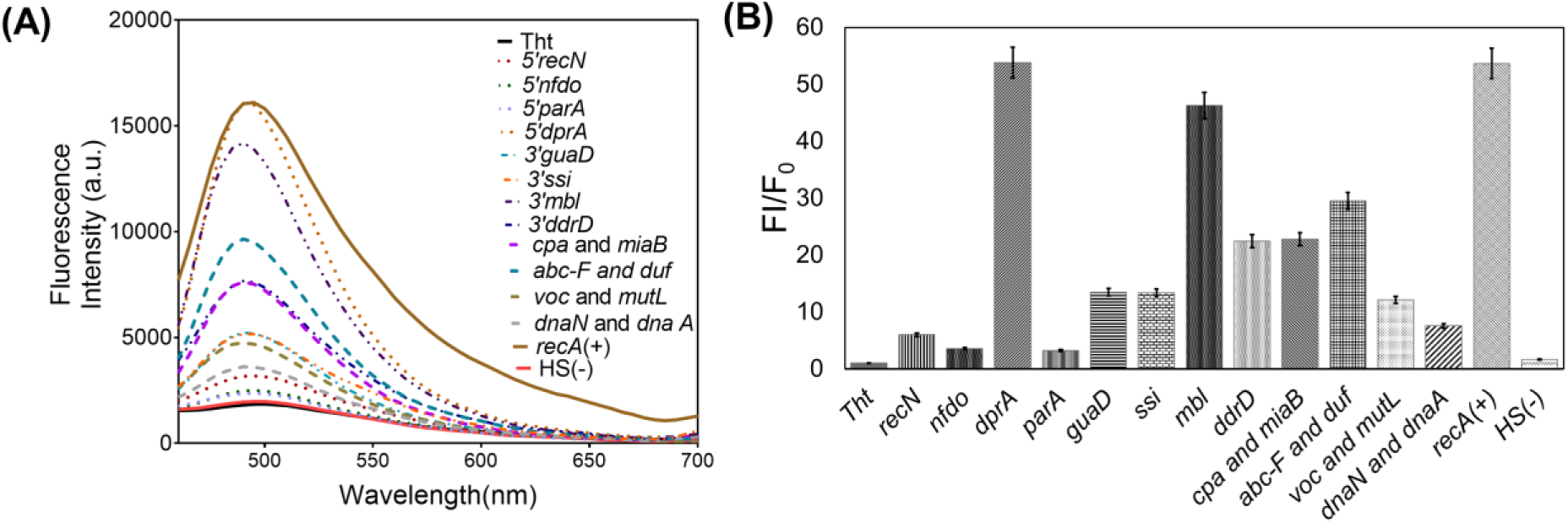
Invitro validation of putative G-quadruplex forming structural motifs using Thioflavin T Fluorescence assay. **(A)** Fluorescence emission spectra of Thioflaviin T in the presence of monovalent cation KCl (100 mM) alone and in the presence of synthesized putative G4s. The fluorescence spectra were collected between 440 and 700nm after excitation at 425 nm**. (B)** The bar graph represents the fold change in fluorescence enhancement of Tht and error bars correspond to 95% confidence intervals. A non-G4 forming sequence (HS) served as a negative control, while *recA* was used as a positive control.

For checking the distinct topologies obtained by G4 sequences with various metal ions, circular dichroism spectroscopy was performed. The circular dichroism spectra of G4 DNA are indicative of their topology. Parallel structures display a maximum absorbance at approximately 264 nm and a minimum at 240 nm, antiparallel structures exhibit a maximum at 290 nm and a minimum at 260 nm, and hybrid structures present a composite profile with maxima at both 290 and 264 nm and a minimum at 240 nm. G4 topologies were checked with monovalent cations like K^+^, Na^+,^ and Li^+^. The CD results revealed that in the presence of KCl, all of the PQS were stabilized and formed parallel topologies irrespective of their positions. The NaCl presence shows distinct topology -parallel (3’-*guaD*; intergenic-*dnaN and dnaA* and *abc-F and dcp*), hybrid (3’-*mbl* and *ssi*, 5’- *recN*,*dprA*and *nfdo*) and anti-parallel (3’-*ddrD* and intergenic-*voc family and mutL* and *cpa and miaB*) topology were observed. All the PQS were destabilized in the presence of LiCl (except intergenic-*cpa and miaB* show anti-parallel topology) (Fig 2A to 2N).

**Figure 2:**
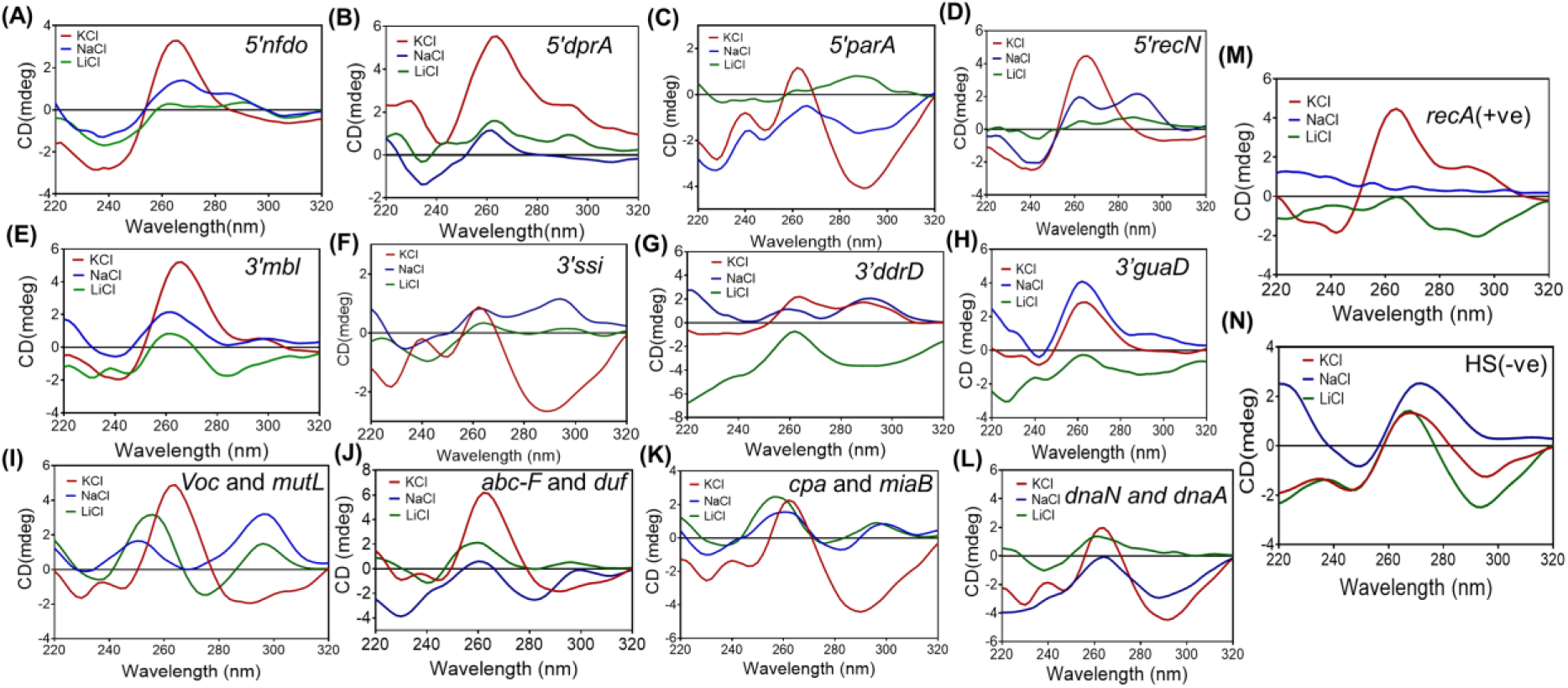
Circular dichroism analysis of putative guanine runs present at various positions (5′/3′/intergenic) in the presence of monovalent cations 100 mM KCl, 50 mM NaCl, and 50 mM LiCl. **(A-D)** Putative G4 motifs present at 5′ end of the gene, **(E-H)** 3′ end of the gene, and **(I-L)** intergenic region. RecA(+) indicates positive control **(M)** and HS is negative control **(N).**

Previous studies also reported the differential effect of monovalent cations on the folding of G4 structures. For example, in the folding kinetics of telomeric G4s studied in the presence of monovalent cations, it was noticed that with K^+^ (50 mM) complete G4 folding was achieved, but with Na^+^ (150 mM), some extent of ssDNA remains (You et al. 2017). In general, Li^+^ cannot support the G4 folding because of its smaller size but can influence/promote the folding fraction differentially with K^+^ or Na^+^. All the putative G4s studied here folded to different topologies depending on K^+^ or Na^+^ present in the solution. Only one putative G4 identified in the intergenic region (*cpa and miaB*) showed stable G4 folding with lithium.

### 3.2 Manganese destabilizes the G4 structures invitro

The folding of PQS in the presence of divalent cations (Mg^2+^ and Mn^2+^) was also checked. With Mg^2+,^ (10 mM) the PGQs were stable (3’-*guaD*, 5’-*recN*, intergenic-*dnaN and dnaA* and *recA*), and formed parallel structures but surprisingly all of them were destabilized when annealed in the presence of 10 mM Mn^2+^ (Fig 3A &3B). Even at lower concentrations of Mn^2+^ (0.1 mM & 2 mM), stable G4s are not formed (Fig 3C & 3D). ThT fluorescence enhancement assay also revealed that in the presence of Mn^2+^ G4s are not stabilized, as an increase in ThT fluorescence was not observed. However the control sequence in KCl presence showed fluorescence enhancement (Fig. 3E).

**Figure 3:**
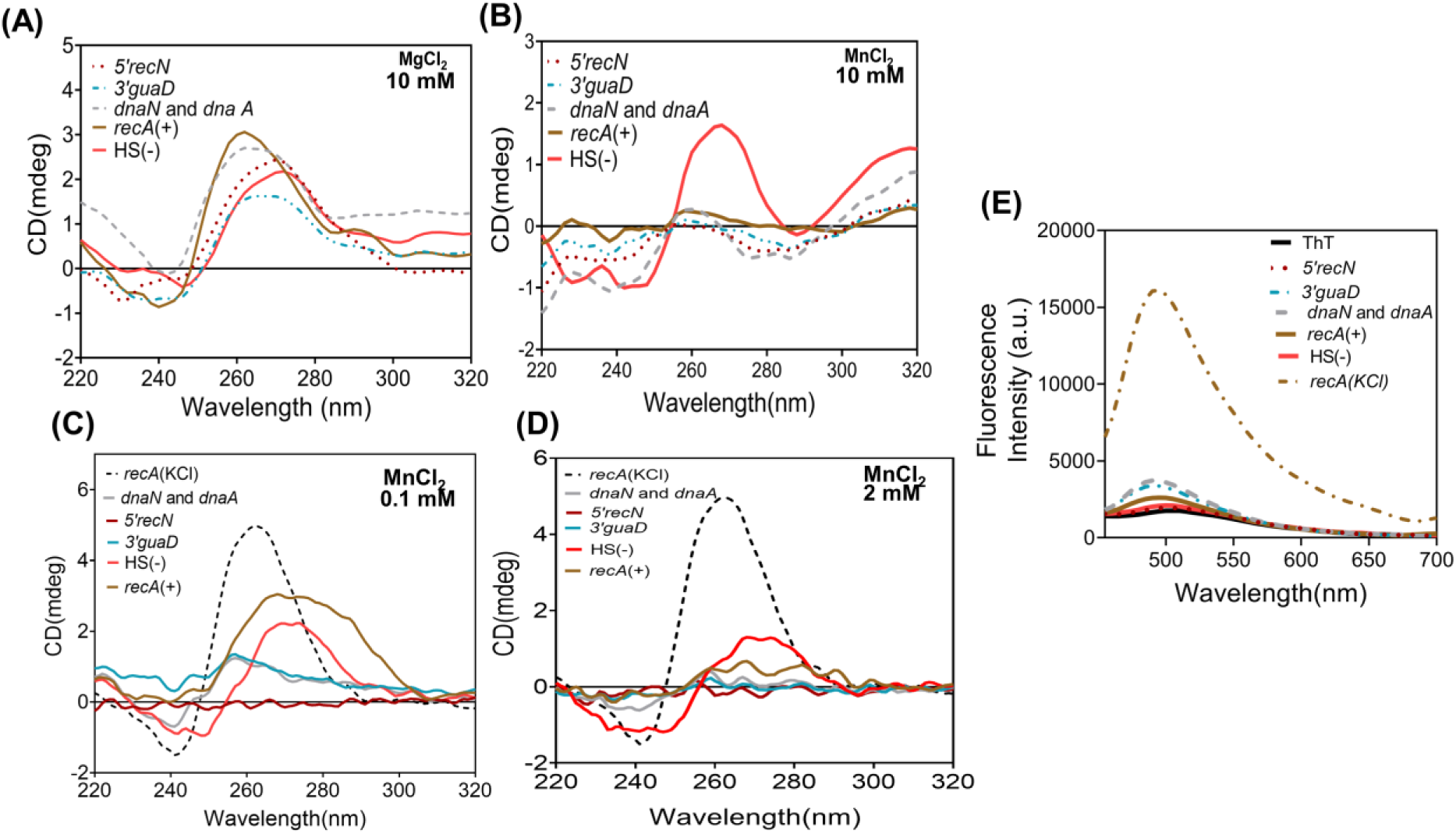
Effect of divalent cation on topology of putative G-quadruplexes. **(A)** CD spectra of synthesized putative G4s in the presence of 10 mM MgCl₂. The solid line indicates the controls. **(B)** CD spectra of synthesized putative G4s in the presence of 10 mM MnCl₂, **(C)** 0.1 mM MnCl_2,_ and **(D)** 2 mM MnCl_2_. **(E)** The fluorescence emission spectra of Thioflavin T in the presence of 10 mM MnCl₂.

In *Deinococcus radiodurans* manganese (Mn^2+^), plays a crucial role in radioresistance. Mn^2+^ forms small-molecule antioxidant complexes, with peptides and phosphate, which protect proteins from oxidative damage caused by ionizing radiation (Culotta VC and Daly MJ; 2013). This study showed that Mn^2+^ (0.1 to 10 mM) could not support the G4 structure formation in vitro by PQS identified on the genome of this bacterium. It is possible that the intracellular levels of Mn^2+^ could interfere with the G4 structure formation under normal growth conditions as cells are not sensitive to G4 ligands under these conditions. But, how Mn^2+^ insmall-molecule antioxidant complexes can modulate G4 structure dynamics is not known.

### 3.3 In vivo G4 structure dynamics in response to cellular stresses

*D. radiodurans* is known to be resistant to other DNA-damaging agents like UV radiation, H_2_O_2,_ and mitomycin C besides gamma radiation. To investigate whether G4 structure dynamics has any role in the cell survival of these DNA-damaging agents or not, growth analysis was performed. Growth studies indicated that in the presence of G4 ligands, the cells showed sensitivity to MMC treatment (5 μM), in addition to gamma treatment, and no change was observed when the cells were subjected to UV and H_2_O_2_ stresses (Fig 4A, 4B, 4C, & 4D). Mitomycin C induces interstrand DNA crosslinks, which can lead to double-strand breaks. This may be the reason this bacterium also showed sensitivity to Mitomycin C treatment in the presence of G4 stabilizers.

**Figure 4:**
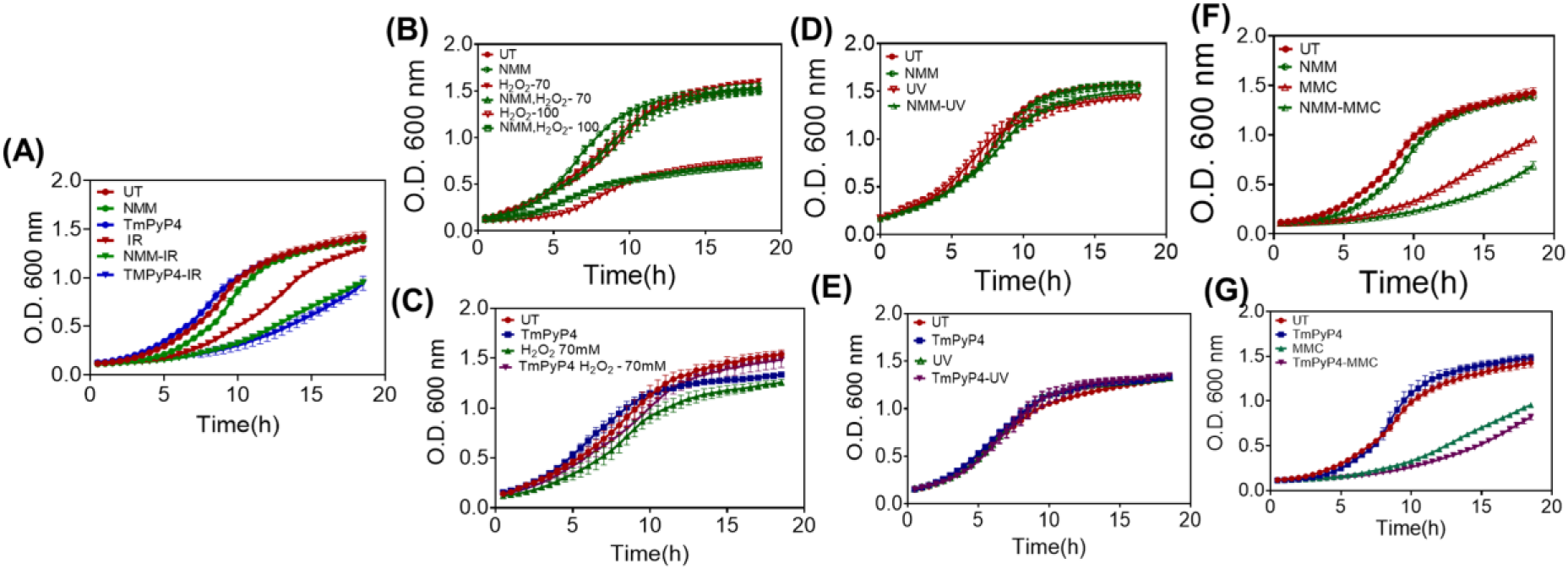
Effect of various DNA damaging agents on cell survival. **(A)** Effect of γ-radiation (6 kGy) in the presence of NMM and TmPyP4 **(B)** Effect of H_2_O_2_ (70 mM & 100 mM) with NMM **(C)** Effect of H_2_O_2_ in the presence of TmPyP4 **(D)** Effect of UV (1000 J/m^2^) in presence of NMM. **(E)** Effect of UV with TmPyP4 **(F)** Effect of MMC (5 µM) with NMM **(G)** Effect of MMC in the presence of TmPyP4.

Next, the in vivo formation of G4 structures in *D. radiodurans* was checked by Thioflavin T(ThT) staining and immunofluorescence experiments as described in the methodology. As shown in Fig. 5A, in the cells that were grown under normal conditions, minimal intensity of ThT was detected suggesting very fast dynamics/ rare formation of these secondary structures. But after gamma radiation treatment at 1h PIR, the intensity of ThT increased and a few prominent high-intense foci/accumulation of ThT were also noticed. At 4h PIR, the mean fluorescent intensity of ThT decreased compared to 1 h PIR (Fig 5B). This suggests during the early period of PIR, a higher number/clustering of G4 structure formation occurs in the cells which may likely resolved due to their dynamic nature at later stages of PIR.

**Figure 5:**
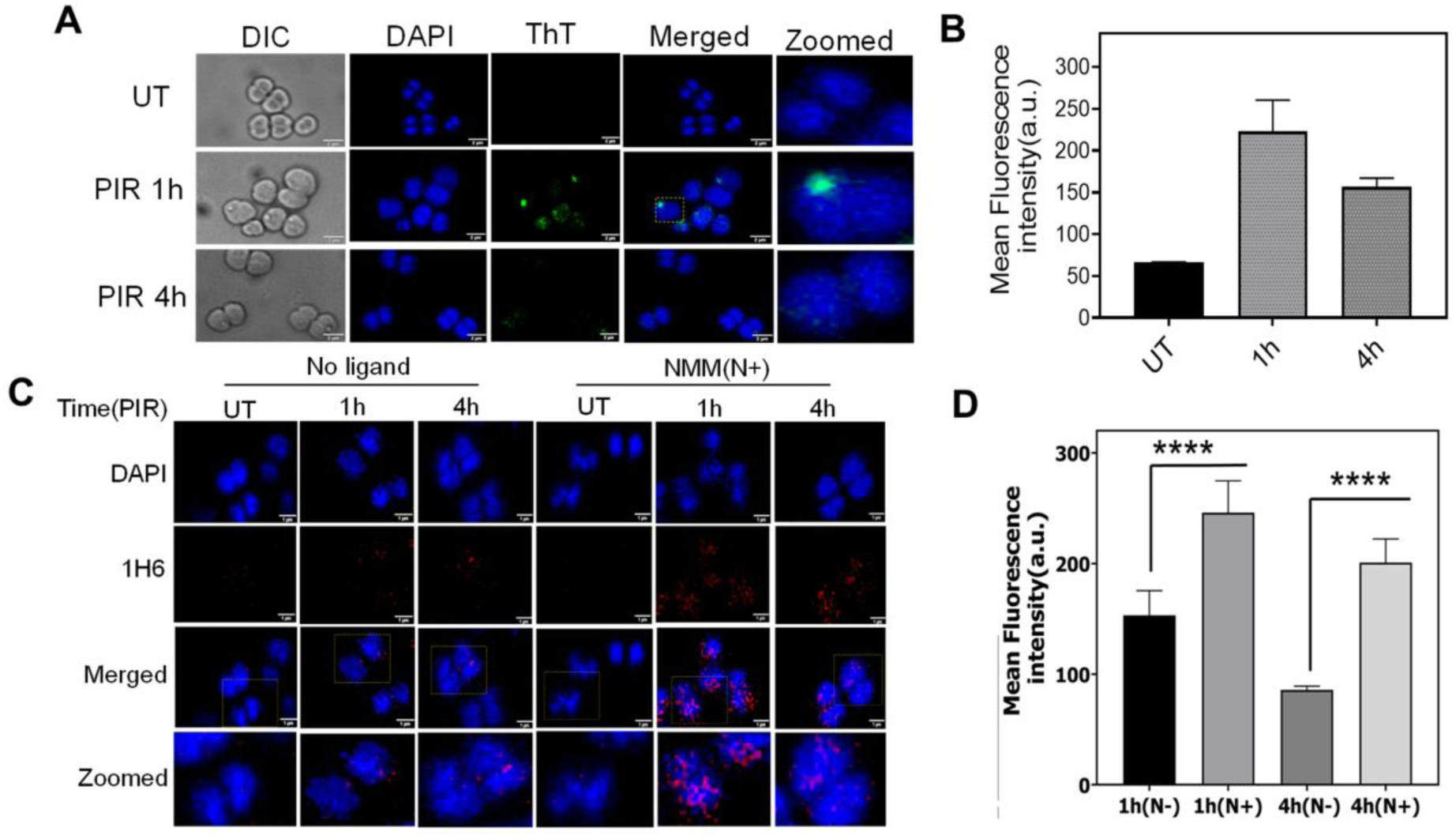
In vivo formation of G4 structure and their dynamics during post-irradiation recovery. **(A)** Fluorescence microscopic examination of *D. radiodurans* cells stained with Thioflavin T after gamma-radiation. At 1h and 4h post-irradiation(PIR) samples were collected and stained with Thioflavin T to visualize the G4 structure and UT represents the unirradiated sample. Images were taken in differential inference contrast (DIC), DAPI (for nucleoid), FITC (for ThT), and merged channels for DAPI and FITC. The scale bar is 2µM. The images shown here are representative pictures of the experiments conducted at least three times. **(B)** The mean fluorescence intensity was calculated using in-built Cell sense software and statistical analysis of post-irradiated samples 1h and 4h were compared to UT cells done in 200–250 cells and plotted. The P-values attained at 95% confidence intervals are depicted as (****) for <0.0001. **(C)** Microscopic images of *D. radiodurans* cells subjected to gamma radiation (6 kGy) in the absence of G4 ligand (No ligand) or presence of G4 ligand [NMM(N+)] and post-radiation recovering cells at 1 h and 4 h (PIR 1 h & 4 h) for visualization of in vivo G4 structure formation using G4 structure-specific antibody (clone 1H6). UT represents the unirradiated cells grown in the absence or presence ofG4 ligand. The microscopic images were taken in different channels, DAPI (for nucleoid), TRITC (for G4 formation), and merged channels for DAPI and TRITC. The zoomed panel represents the image of individual cells for better visualization (scale bar is 1µm). The representative images are of biological triplicates. **(D)** The mean fluorescence intensity was calculated using automated cell sens software and graphs were plotted using GraphPad Prism and statistical analysis was done in 100-150 cells using the student’s t-test. The p-values obtained at 95% confidence intervals are indicated as (****) for <0.0001 and ns(non-significant) >0.05.

Further, the formation of G4 structure in vivo, and the effect of G4 binding ligands on their dynamics during PIR was also monitored using an anti-DNA G-quadruplex (G4) antibody as described earlier. NMM-treated and untreated samples subjected to gamma radiation were checked for in vivo G4 structures (Fig 5C). The microscopic results indicate that 1 h PIR samples have more fluorescence signal compared to untreated samples (UT) indicating in vivo formation of G4 structure after gamma radiation treatment. The presence of NMM during post-irradiation further increased the fluorescence signal (1 h NMM) suggesting cells have more stabilized G4 structures after G4 ligand treatment. At 4 h PIR also NMM treated samples showed a higher signal than ligand untreated samples (Fig.5D).

The formation of G4s in cells treated with other stresses was also observed. Results revealed small dispersed foci of ThT at 1h in cells treated with UV and H_2_O_2_, and the number of such foci reduced at 4 h. In response to MMC treatment, the number of dispersed foci was observed at 4 h (Fig. 6). Nevertheless, under all the observed stresses, cells at 1 h PIR condition showed more intense /accumulation of ThT foci. Earlier G4 structure role in radioprotection across different regions of the human genome leading to differential radiosensitivity was also demonstrated (Kumari et al. 2019; Kumari and Raghavan 2021). Further, stress also induces stable RNA quadruplex folding, which increases their stability (Kharel et al. 2023). Similarly in *D. radiodurans* the G4s formed may assist in radioprotection or may be involved in protein expression/recruitment to repair the damaged DNA after exposure to DNA-damaging agents.

**Figure 6:**
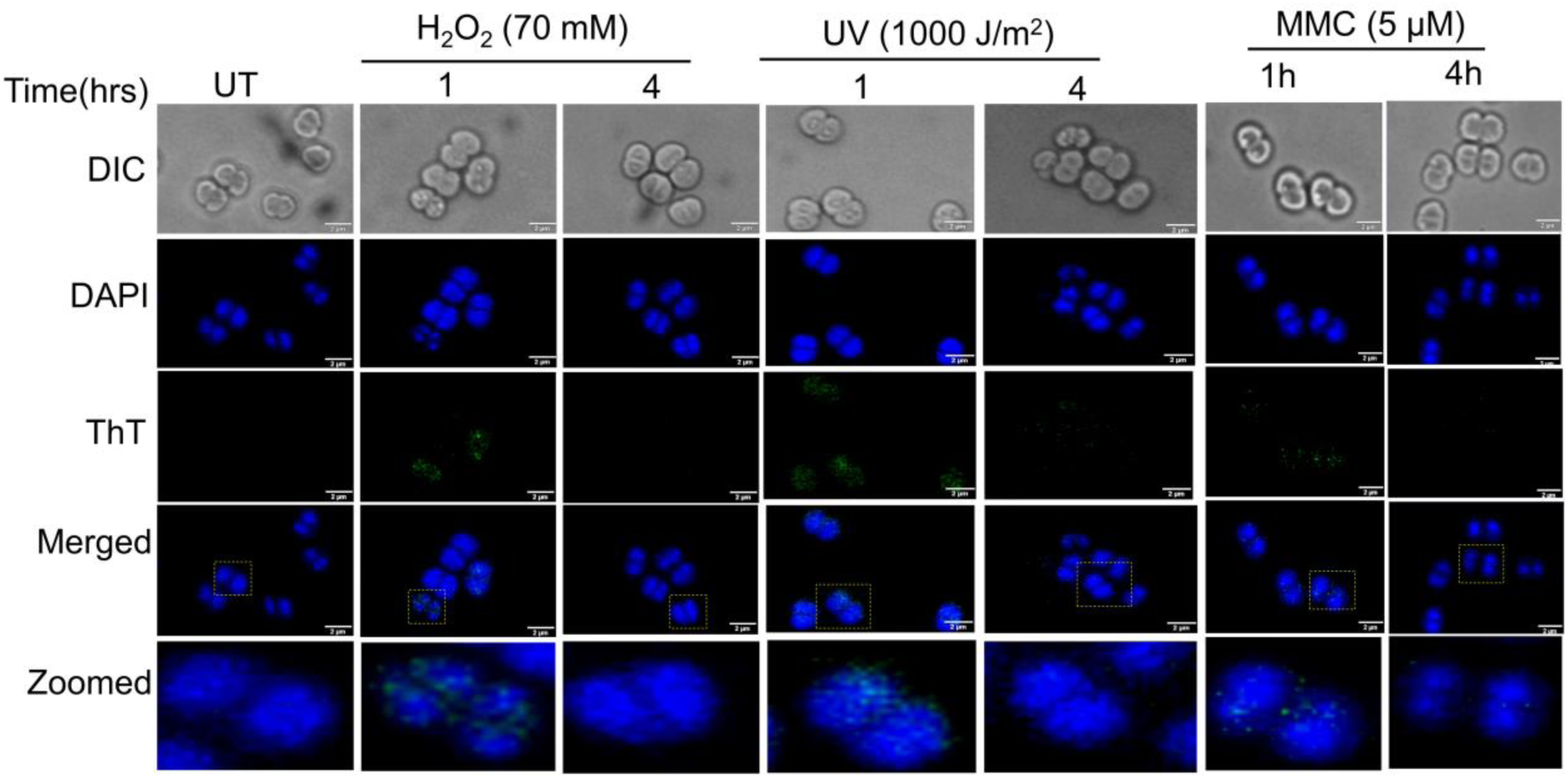
Microscopic images showing the in vivo formation of G4 structure under different DNA damaging agents. **(A)** Wild-type cells (R1) were treated with H_2_O_2_ (70 mM), UV (1000 J/m^2^), and MMC (5 µM) **(B)** UT represents the untreated samples. Images were taken in differential inference contrast (DIC), DAPI (nucleoid), FITC (ThT), and merged (DAPI+FITC) channels and are presented in planar view. The scale bar for all the panels is 2µm.

### 3.4 Arrest of G4 structural dynamics slows DSB repair kinetics

*D. radiodurans* cells showed more number of G4 structures at 1 h PIR, while in the presence of NMM, further increment was noticed. To repair DSBs mainly ESDSA pathway operates in this bacterium. PFGE is generally used to follow DNA damage repair kinetics. Hence, PFGE was performed with *D. radiodurans* cells that were collected at different time intervals after gamma radiation treatment. The data analysis showed that NMM-treated cells exhibited slower DSB repair kinetics compared to the wild type. In untreated cells (NMM -), the recovery of DNA bands was started after 2 h of post-irradiation, while in NMM treated cells (NMM +) the band appearance started around 4 h of PIR suggesting delayed repair in cells that were treated with G4 ligands (Fig. 7A). The results revealed that the stabilization of guanine quadruplex slows the repair kinetics making the cells sensitive to G4 binding drugs after gamma radiation after post-irradiation recovery in *D.radiodurans*.

**Figure 7:**
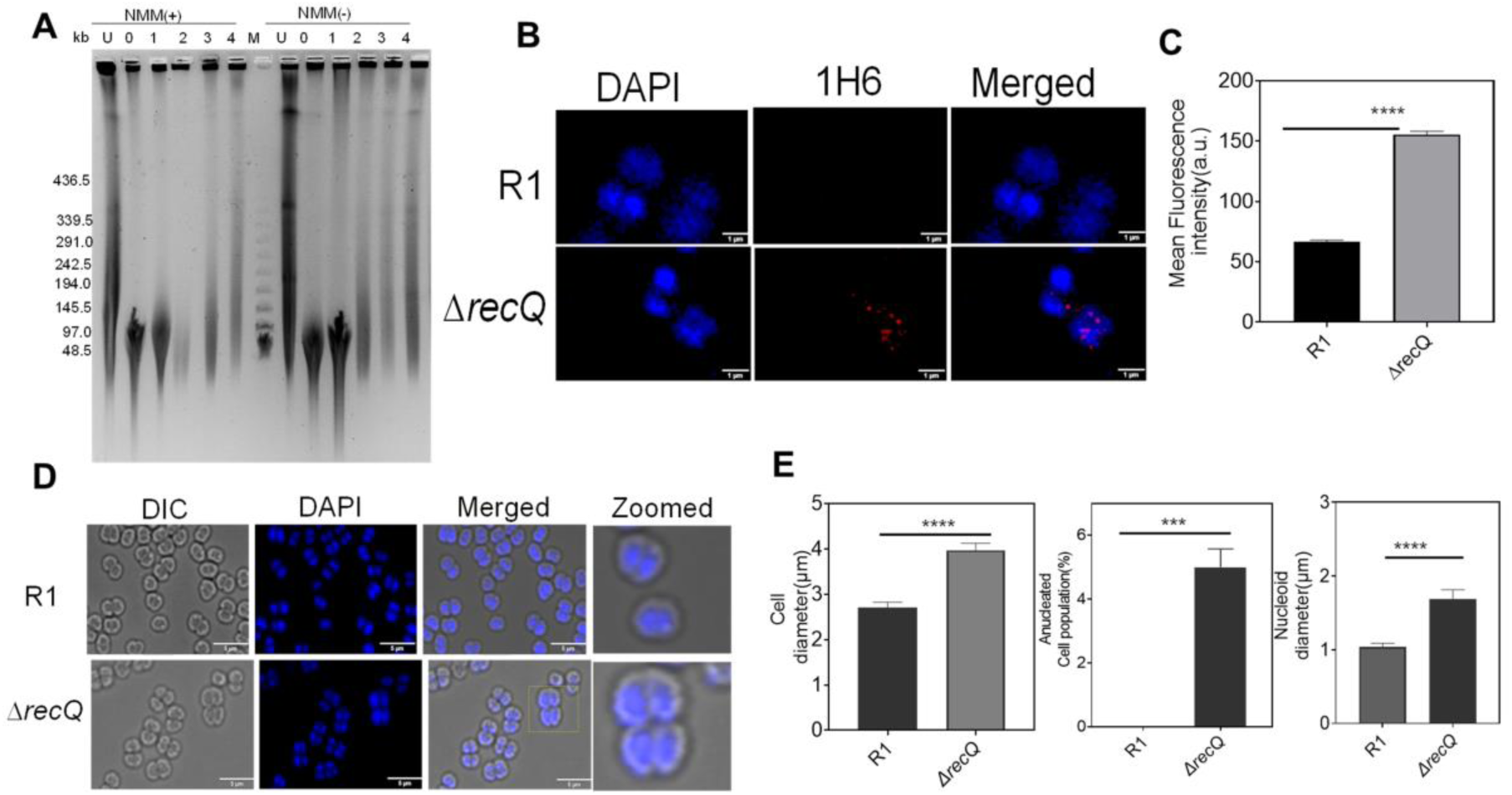
Effect of stabilization of guanine quadruplex structures on double-strand break repair and genome stability in *D. radiodurans.* **(A)** *Deinococcus* cells were grown in the presence (NMM+) and absence (NMM-) of the G4 binding ligand. Gamma radiation untreated (UT) and post-irradiation recovering cells at different time intervals 0 h, 1 h, 2 h, and 4 h were subjected to PFGE along with Lambda PFG marker (M) to check the double-strand break repair kinetics. (**B)** Immunofluorescence assay using an anti-G4 antibody (clone1H6) of wild-type(R1) and *recQ* mutant (*ΔrecQ*). These cells were observed under DAPI (for nucleoid) and TRITC (1H6) channels. The merged images represent the DAPI+TRITC channel. **(C)** The mean fluorescence intensity was depicted for R1 and *ΔrecQ.* Data were analyzed using Student’s t-test (**** P < 0.0001) and the data was plotted using GraphPad Prism software 8. Morphological alterations in Δ*recQ* mutants compared to wild-type (R1). **(D)** Microscopic images show the morphological changes observed in *ΔrecQ* mutants compared to wild-type (R1) under the microscope. DIC, DAPI, and merged channels are shown. The scale bar is 5µm. **(E)** The cell size distribution, Anucleated cell population, and the nucleoid size distribution. The GraphPad Prism software 8 was used for statistical analysis of the different morphologies. The p-values attained at 95% confidence intervals are depicted as (***) for <0.001 and (****) for <0.0001. Each experiment was repeated thrice and biological replicates were included.

### 3.5 RecQ helicase is involved in the regulation of G4 structural changes and genome maintenance

In *D. radiodurans,* RecQ plays an important role in the metabolism of G4 structures in this bacterium (Khairnar et al. 2019). Hence, the in vivo G4 structure formation in Δ*recQ* under normal conditions was examined by microscope studies. The results reveal that *ΔrecQ* shows more G4 structures even under normal conditions (*ΔrecQ* UT vs R1 UT, Fig. 7B). Further, to check the effect of an increase in G4s on genome maintenance, various parameters like cell size, nucleoid diameter, and the percent of anucleated cells in a population (∼105-200 cells) compared to wild type. As shown in the figure, a statistically significant change in the average size of the cells (fold change-1.461, P-value <0.0001), the diameter of the nucleoids (fold change-1.631, P-value <0.0001), and anucleated cell population (P-value <0.001) were observed in *ΔrecQ* cells compared to wild-type (Fig. 7C). All these results suggests that RecQ helicase is the main protein involved in the regulation of G4 structure folding and unfolding and genome maintenance in *D. radiodurans*.

The association of G4 structures in cellular processes like transcription, DNA replication, DNA damage repair, epigenetics, etc. are well studied in eukaryotes. G4-forming sequences were also identified in bacterial genomes, where these sequences are non-randomly distributed, and predominantly associated with gene regulatory regions (Cueny et al., 2022). But unlike in eukaryotes, there exist fewer studies that probed the in vivo role of guanine quadruplexes in bacteria. This study has first time shown the in vivo G4s formation in response to DNA-damaging agents. Numerous putative G4 motifs are located in the genome of *Deinococcus radiodurans.* Under normal growth conditions, detectable levels of G4s are not formed. This may be due to the intracellular concentration of Mn^2+^ or the presence of G4 helicases that may disrupt/resolve the G4s. When exposed to stress, the G4s increase in cells, especially to gamma radiation. These G4s formed may regulate the transcription of genes or DNA repair by recruitment of different proteins. The arrest of G4 structure formation/resolution by G4 ligands like NMM may result in stable secondary structures that interfere with the DNA repair process causing sensitive phenotype. At the same time, accumulations of G4s during normal growth conditions as seen in *ΔrecQ* cells may impede the cellular processes leading to genome instability. Thus, during the evolution extremophiles like *D. radiodurans* might have utilized the guanine quadruplexes to combat the cellular stresses. Further studies on regulatory pathways/proteins that act to maintain the G4s frequency in this bacterium under different cellular conditions are crucial to understanding the G4 biology in prokaryotes.

## Acknowledgment

We sincerely thank Dr. Hema Rajaram, Bhabha Atomic Research Centre, Mumbai, for her suggestions and support while pursuing this work. Ms. Himani Tewari expressed her thanks to the Department of Atomic Energy, Government of India for the research fellowship. We also thank Ms. Sindhuja Singh for her input during the manuscript preparation.

## Authors’ contributions

HT performed the experiments, analyzed the data, and wrote the manuscript. SM analyzed the data and wrote the manuscript. SK conceptualized the idea, analyzed the data, and wrote the manuscript.

## Conflict of interest statement

The authors declare no conflict of interest.

## Notes

### Competing Interest Statement

The authors have declared no competing interest.

